# RRNPP_detector: a tool to detect RRNPP quorum sensing systems in chromosomes, plasmids and phages of gram-positive bacteria

**DOI:** 10.1101/2021.08.18.456871

**Authors:** Charles Bernard, Yanyan Li, Eric Bapteste, Philippe Lopez

## Abstract

Gram-positive bacteria (e.g. Firmicutes) and their mobile genetic elements (plasmids, bacteriophages) encode peptide-based quorum sensing systems (QSSs) that regulate behavioral transitions in a density-dependent manner. In their simplest form, termed “RRNPP”, these QSSs are composed of two adjacent genes: a communication propeptide and its cognate intracellular receptor. Despite the prime importance of RRNPP QSSs in the regulation of key biological pathways such as virulence, sporulation or biofilm formation in bacteria, conjugation in plasmids or lysogeny in temperate bacteriophages, no tools exist to predict their presence in target genomes/mobilomes. Here, we introduce RRNPP_detector, a software to predict RRNPP QSSs in chromosomes, plasmids and bacteriophages of gram-positive bacteria, available at https://github.com/TeamAIRE/RRNPP_detector. RRNPP_detector does not rely on homology searches but on a signature of multiple criteria, which are common between distinct families of experimentally-validated RRNPP QSSs. Because this signature is generic while specific to the canonical mechanism of RRNPP quorum sensing, it enables the discovery of novel RRNPP QSSs and thus of novel “languages” of biocommunication. Applying RRNPP_detector against complete genomes of viruses and Firmicutes available on the NCBI, we report a potential 7.5-fold expansion of RRNPP QSS diversity, alternative secretion-modes for certain candidate QSS propeptides, ‘bilingual’ bacteriophages and plasmids, as well as predicted chromosomal and plasmidic Biosynthetic-Gene-Clusters regulated by QSSs.

## INTRODUCTION

Quorum sensing is the mechanism by which microbial entities sense when their population density reaches a threshold level, and thereupon typically switch from individual to group behaviors (1). The population density is reflected by the extracellular concentration of a communication signal, produced and secreted by individual entities. The quorum is met when this signal reaches a threshold concentration, at which it starts to be robustly detected and transduced population-wide by its cognate receptor module. In bacteria of the Proteobacteria or Actinobacteria phyla, these communication signals typically are small molecules synthesized by enzymes (2,3) whereas in the Firmicutes phylum, these are oligopeptides, matured from genetically encoded pro-peptides (4). Peptide-based quorum sensing systems can be divided into two main categories: those with a receptor module composed of a membrane-bound sensor coupled with an intracellular response regulator (two-components system) (5), and those in which the receptor is an intracellular transcription factor (or a protein inhibitor) that gets either turned-on or -off upon binding with the imported communication peptide (one component system) (6,7). The latter are generally included under the term RRNPP, named after the 5 first experimentally-characterized families of such receptors: Rap (*Bacillus* genus), Rgg (*Streptococcus* genus), NprR (*Bacillus cereus* group), PlcR (*Bacillus cereus* group) and PrgX (pCF10 plasmid of *Enterococcus faecalis)* (7–9).

The initial members of the RRNPP group of QSSs were reported to trigger key biological pathways when their encoding population reach high densities: from virulence (Rgg, PlcR) to competence (Rgg, Rap), necrotropism (NprR), sporulation, biofilm formation (Rap, NprR) and inhibition of conjugation (PrgX) (7–9). Considering that the virulence of *Bacillus* and *Streptococcus* pathogens may cause infectious diseases in humans (10,11), that the spore is the transmissive form of many *Bacillus* and *Clostridium* human pathogens (12), that biofilms contribute to infections or food-poisining (13–15), and that competence and conjugation are responsible for the spread of antibiotic resistance genes (16), RRNPP QSSs are directly linked to central health issues.

Interestingly, the case of the plasmidic PrgX system illustrates that RRNPP QSSs may not only be present on bacterial chromosomes but also on mobile genetic elements (MGEs). Consistently, the conjugation-regulating TraA-Ipd1 system have later on expanded the list of plasmidic RRNPP QSSs (17). In 2017, Erez *et al*. even discovered that some temperate bacteriophages encode RRNPP QSSs, namely the “arbitrium” systems that guide the lysis-lyogeny decision upon *Bacillus* infection (18,19). Recently, two other chromosomal RRNPP QSSs have been experimentally validated: the QsrB-QspB system that delays sporulation and solvent formation in *Clostridium acetobutylicum* (20) and the AloR13-AloP13 sporulation-regulator in *Paenibacillus polymyxa* (21). As important the experimentally-validated RRNPP QSSs are in the regulation of the biology of their encoding microbial entity, they represent only the tip of the iceberg. Indeed, the diversity of RRNPP QSSs is likely not fully explored, as hinted by the numerous candidate RRNPP QSS receptors in *Clostridium* and *Enterococcus* species reported to harbor local similarities with regions of Rap, NprR, PlcR or Rgg (20,22,23). Uncovering novel RRNPP QSS families would unveil new bacterium-bacterium, plasmid-plasmid or phage-phage modes of communication but most and foremost reveal novel density-dependent evolutionary strategies that could transform our views of microbial interaction, adaptation and evolution. Expanding RRNPP QSS diversity would also likely pave way to major practical outcomes as novel RRNPP QSSs could regulate the production of novel antimicrobial compounds (24,25) or could underlie mechanisms by which some human pathogens acquire virulence (26–29).

Conveniently, we noticed that the different members of the RRNPP group share a common signature of 5 criteria (**Fig. 1**): i) the pro-peptide is a small protein (10-70aa), ii) the pro-peptide is secreted, except in rare exceptions (e.g. Shp and PrgQ (7)), via the SEC-translocon and further matured by exopeptidases into a communication peptide, iii) the receptor has a length comprised between 250 and 500aa, iv) the receptor harbors tetratricopeptide repeats (TPR), which are structural motifs involved in the binding of small peptides (in this case, the cognate communication peptide), v) the genes encoding the pro-peptide and the receptors are directly adjacent to each other. Advantageously, a large amount of reference Hidden-Markov-Models (HMMs) from the Cath-Gene3D (30), Superfamily (31), SMART (32), Pfam (33) and TIGRFAM (34) databases are already available to detect TPRs in protein sequences. Moreover, a tool called SignalP specifically computes the likelihood that proteins harbor a signal sequence for the SEC-translocon (35) (**Fig .1**). Consequently, the generic, yet specific signature of RRNPP QSSs is detectable *in silico*, without requiring homology searches that would limit the output to representatives of already known QSSs. On this basis, we have developed RRNPP_detector, a python software that detects the RRNPP signature in chromosomes and MGEs of gram-positive bacteria. We present its workflow, its usage and showcase practical examples of analyses against the complete genomes of viruses and of Firmicutes available on the NCBI.

**Figure 1:**
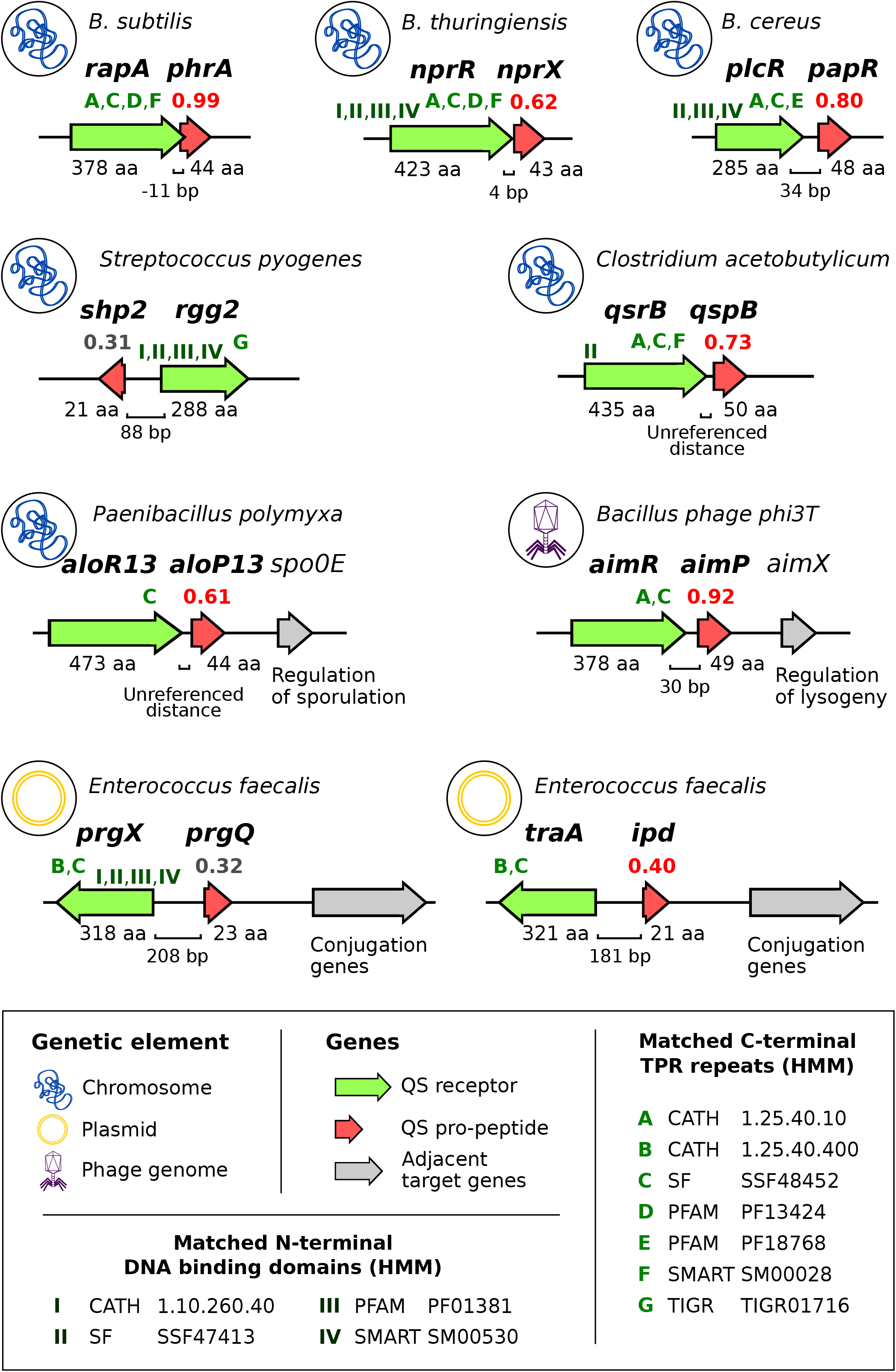
Common features between experimentally validated RRNPP-type QSSs. Each genomic context corresponds to the representative QSS of an experimentally characterized RRNPP-type QSS family. The icon at the left of each context indicates the genetic element that encodes the QSS (bacterial chromosome, phage genome or plasmid) and the associated label indicates the taxon to which this genetic element belongs. The green gene corresponds to the quorum sensing receptor and the red gene to its cognate propeptide, whereas a grey gene indicates an adjacent, target gene (or set of genes) demonstrated to be regulated by the QSS. The intergenic distance between the receptor and the propeptide genes is given in number of base pairs and the length of receptors and propeptides is given in number of aminoacids. The number above each pro-peptide corresponds to the likelihood, computed by SignalP, that the propeptide harbors a N-terminal signal sequence for the SEC-translocon. A likelihood score colored in red means that the propeptide is predicted by SignalP to be secreted via the SEC-translocon, whereas a score colored in grey means that it is predicted to be secreted otherwise. Shp2 and PrgQ are the only two propeptides predicted to be secreted otherwise, consistent with the fact that they are the only ones known to be secreted via another secretion system than Sec(SEC/SpI), namely the PptAB complex (7). The green letters above the C-terminal encoding region of each receptor indicate the names of the HMM (PFAM, SMART, TIGR) or of the HMM family (CATH, SuperFamily) of Tetratricopeptide repeats (TPRs) that are found within the sequence of the translated protein. The roman numbers above the N-terminal encoding region of each receptor indicate the names of the HMM or of the HMM family of DNA binding domains found in the sequence of the translated protein.

## MATERIALS AND METHODS

### Overview of the RRNPP_detector workflow

Given a fasta of protein sequences and the coordinates of their coding sequences in target genome(s), Metagenomics-Assembled-Genome(s) or contig(s), RRNPP_detector first reduces the search space by retaining only proteins that have a length compatible with RRNPP receptors (250-500aa) and pro-peptides (10-70aa). Next, only the 250-500aa proteins whose coding sequences (CDSs) are found adjacent with the CDS of a 10-70aa protein are retained, and vice versa. Then, HMMs of TPRs found in reference RRNPP receptors (PFAM: PF13181, PF13424; Gene 3D: 1.25.40.400, 1.25.40.10; TIGRFAM: TIGR01716; Superfamily: 48452, SMART: SM00028) are queried with hmmsearch (36) against the remaining 250-500aa proteins. Only the proteins matched by at least one HMM of TPRs (E-value < 1E-5) are retained and designated as potential receptors. These potential receptors next undergo a computational characterization. First, a BLASTp search (37) of reference RRNPP receptors (Rap, Rgg, NprR, PlcR, PrgX, TraA, AimR of Bacillus phage phi3T, AimR of Bacillus phage Waukesha, QsrB and AloR13) against these potential receptors are launched to identify the homologs of already known QSSs (by default: E-value <= 1E-5, identity >= 20%, and alignment coverage >= 60% of the length of both the query and the target sequences). Then, an HMMsearch of DNA-binding domains found in reference receptors (PFAM: PF01381; Gene3D: 1.10.260.40; Superfamily: 47413; SMART: SM00530) is launched against the potential receptors to identify those that may be transcription factors. After this characterization step, the 10-70aa proteins are further reduced to those with a CDS adjacent to the one of a potential receptor. At this stage, each tandem of remaining adjacent pro-peptide and receptor are considered forming a potential QSS and will eventually be outputed. However, either a ‘permissive’ or a ‘conservative’ label will be assigned to these candidate QSSs according to the next filters. First, any QSS with an intergenic distance > 400 base pairs between the two CDSs will be considered as ‘permissive’ because low intergenic distance tends to correlate with functional association (38). Second, only the remaining QSSs with a pro-peptide predicted by SignalP (35) to harbor a signal for the SEC-translocon under the ‘-org gram+’ option will be considered as ‘conservative ’ (**Fig. 2**). Because a few experimentally-characterized pro-peptides are known to be secreted via other systems than the SEC-translocon (*e.g*. Shp2 and PrgQ (7)), we thought that it would be relevant to keep the QSSs in which the pro-peptides are not predicted to be secreted via the SEC-translocon in the ‘permissive’ output.

**Figure 2:**
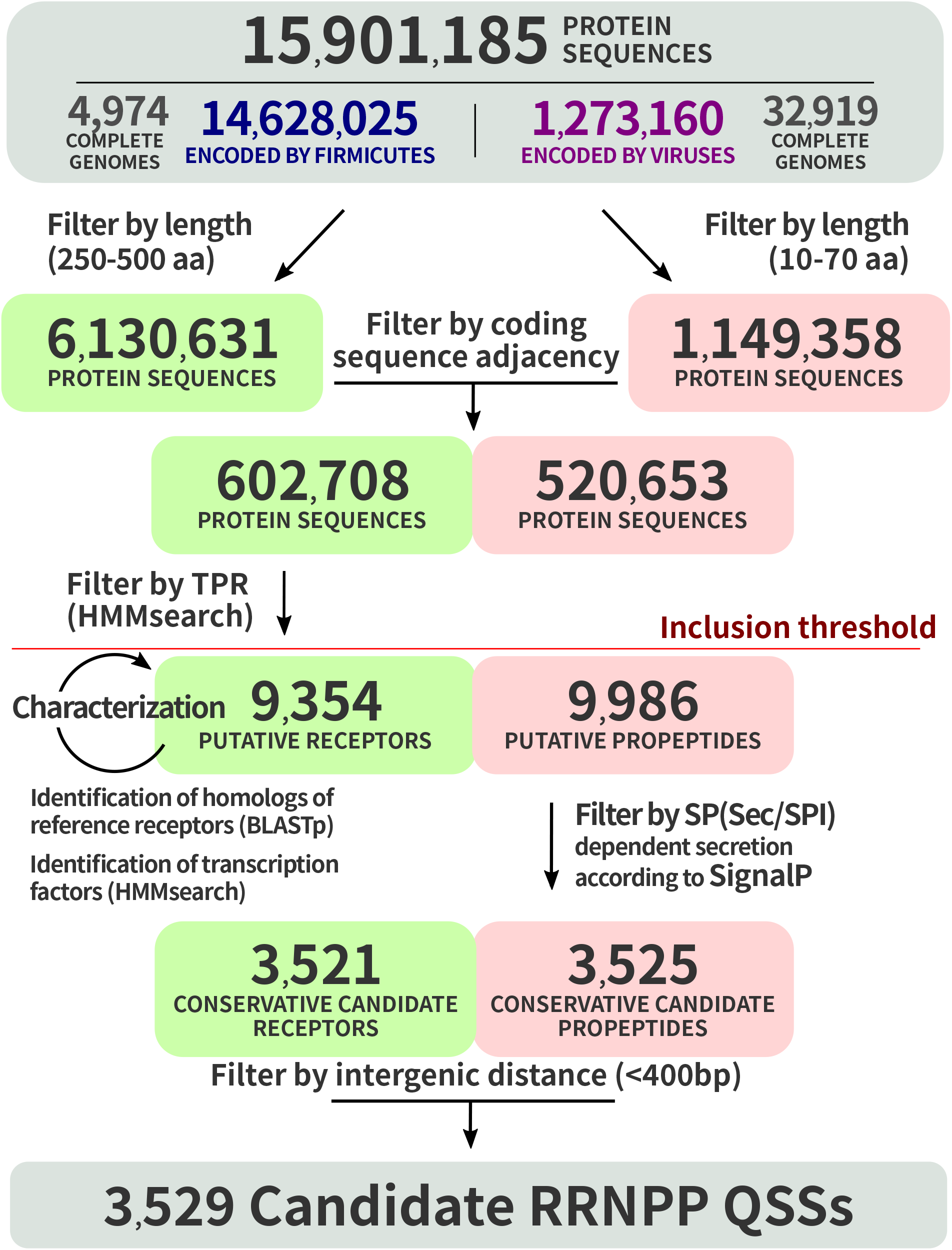
Workflow of RRNPP_detector illustrated with real data from complete genomes of Firmicutes and viruses. RRNPP_detector defines candidate RRNPP-type quorum sensing systems as tandems of adjacent CDSs encoding a candidate receptor (250-500aa protein matching HMMs of peptide-binding tetraticopeptide repeats (TPRs)) and a candidate pro-peptide (10-70aa protein predicted to be excreted via the SEC-translocon). Each green and red rectangle represents a step towards the final identification of candidate receptors and candidate propeptides, respectively. Each arrow represents a filtering operation to narrow down the search space to ‘conservative’ candidate RRNPP QSSs. The horizontal red line indicates the inclusion threshold after which all remaining pairs of adjacent receptors and propeptides will be outputed: pairs that pass all the filters below the inclusion threshold will be considered as ‘conservative’ candidate RRNPP QSSs, whereas the other pairs will be considered as ‘permissive’ candidates. RRNPP_detector identified 3529 ‘conservative’ candidate RRNPP QSSs from 3521 candidate receptors and 3525 candidate propeptides because a receptor can be flanked on both side by a candidate propeptide and vice versa.

### Usage, input and output of RRNPP_detector

RRNPP_detector was designed to take two kinds of input. The first option is to provide a nucleotide sequence (fna) of either one or multiple genetic elements. RRNPP_detector will then call Prodigal (39) to detect the CDSs in these nucleotide sequences and produce a fasta of the predicted protein sequences (faa) and an annotation file of the coordinates of their CDSs (gff). Alternatively, the user can submit a fasta of the protein sequences from one or multiple genomes/MAGs/contigs (faa) along with an annotation file of these genetic elements (in one concatenated gff or in one concatenated NCBI Assembly feature table). Option 2 has the advantage to integrate pre-computed annotations and therefore to keep reference IDs of proteins along the analysis. RRNPP_detector was designed to be launched in two modes: a default fast mode that processes all the target genomes/MAGs/contigs together in a single run; and a ‘preserve_ram’ mode designed to minimize RAM usage at the expense of speed by processing multiple genomes one by one in a for loop. RRNPP_detector outputs a fasta directory with the protein sequences of the ‘permissive’ pro-peptides, the ‘permissive’ receptors, the ‘conservative’ pro-peptides and the ‘conservative’ receptors. In addition, two ‘permissive’ and ‘conservative’ summary tabular files with various information regarding each candidate QSS are produced. A summary file typically displays, for each QSS, the ID of the encoding genetic element, the intergenic distance between the pro-peptide and the receptor genes, the genomic orientation of these two genes, the IDs and the genomic coordinates of the receptor and the propeptide, the best HMM of TPRs and DNA-binding domains found in the sequence of the receptor (with associated E-value), the BLASTp results in case of an homology of the candidate receptor with a reference receptor, and the SignalP prediction of the secretion mode of the pro-peptide (with associated likelihood score).

## RESULTS

### RRNPP_detector reveals thousands of candidate RRNPP QSSs in complete bacterial chromosomes, plasmids and phage genomes

To illustrate RRNPP_detector, we launched it against the 4974 complete genomes of Firmicutes and the 32919 complete genomes of Viruses Firmicutes available on the NCBI Assembly Database (**Fig. 2**). We describe how such analyses can be easily done with the practical example of Viruses in the readme file of RRNPP_detector: https://github.com/TeamAIRE/RRNPP_detector/readme.md. We report the identification of 6467 ‘permissive’ candidate RRNPP QSSs, distributed across 1074 different taxa (**Table S1**). Some receptors of these ‘permissive’ candidate QSSs notably include the RopB regulator of virulence in *Streptococcus pyogenes* (27), as well as additional members of the AimR, AloR13, NprR, PlcR, PrgX, QsrB, RapA and Rgg2 experimentally-validated families that did not pass the ‘conservative’ thresholds of RRNPP_detector (**Table S1**). Interestingly, 105 ‘permissive’ RRNPP QSSs have been detected in 97 isolated genomes of viruses, of which 89 bacteriophages and 8 giant eukaryotic viruses of the *Mimiviridae* taxonomic family. Accordingly, some bacteriophage genomes are found to encode multiple ‘permissive’ candidate QSSs (up to 4 in Brevibacillus phage SecTim467) (**Table S1**). Last but not least, 435 and 47 ‘permissive’ candidate QSSs include a propeptide predicted to be secreted via the LIPO (Sec/SpII) and TAT (TaT/SPI) secretion systems, respectively, pointing at alternative secretion modes than SEC (Sec/SpI) for some candidate RRNPP propeptides (**Table S1**). In addition, we report the identification of 3529 ‘conservative’ candidate RRNPP QSSs that we hereafter characterize in more depth (**Table S2**). We first used the Phaster API (40) for each chromosome or plasmid carrying at least one QSS to identify whether the genomic coordinates of the QSS(s) fell within prophage regions (genomes of lysogenic phages inserted within host genomes) and should be thus considered as viral instead of bacterial. Of the 3529 conservative candidate RRNPP QSSs, we found that 2793 are chromosomal (on 941 different chromosomes), 222 are plasmidic (on 172 different plasmids), 505 are predicted to be prophage-encoded (on 219 predicted intact prophages, 102 questionable prophages and 176 incomplete prophages) and 9 are encoded by genomes isolated from free temperate bacteriophages (on 8 distinct bacteriophages) (**Table S2**). Although the presence of multiple QSSs on single chromosomes is not rare (8,21) due to the selective pressure that may exist for the acquisition of additional subpopulation-specific QSSs (41,42), only 1 MGE (Bacillus phage phi3T) has been previously reported to encode two RRNPP QSSs (43). Here, we report the identification of 30 plasmids and 8 (pro)phages encoding multiple candidate RRNPP QSSs (up to 5 in the pBMB400 plasmid of *Bacillus thuringiensis* serovar kurstaki str. YBR-1520) (**Table S3**). Interestingly, multiple QSSs might enable a MGE to produce distinct communication peptides that accumulate differentially in the medium, hence to sense and react to multiple population density levels (43).

### RRNPP_detector massively expands RRNPP QSS diversity

In order to classify the 3529 conservative candidate RRNPP QSSs, the sequences of the candidate receptors were BLASTed against each other to identify groups of homologs, defined as connected components in the resulting sequence similarity network (wherein two nodes (proteins) are connected if they show a sequence identity >= 30% over more than 80% of the lengths of both sequences, as in (44)) (**Fig. 3**). We report the identification of 31 groups of homologous receptors (with at least 2 homologous sequences in it) and of 29 singletons (any receptor with no clear homology to others) (**Table S2** and **Fig. 3**). If a singleton or at least one sequence in a group of homologous receptors was matched by an experimentally-validated receptor in a BLASTp search (same thresholds as above), we considered this group as already known. Of the 60 different types of receptors detected, 8 were hence considered as already known: the Rap + NprR group (N=2991), PlcR (N=279), AimR of *B. subtilis* phages (N=20), AloR13 (N=16), AimR of *B. cereus* phages (N=10), TraA (N=5), Rgg2 (N=4) and QsrB (N=2) (**Fig. 3** and **Fig. 4**)). Our results thus increase the diversity of RRNPP QSSs by a factor of 7.5 and paves the way to the identification of multiple novel peptide-based biocommunication “languages” (**Fig. 4**). Importantly, these different groups of QSSs cover a substantial phylogenetic diversity. Indeed, they are distributed across 21 bacterial taxonomic families (**Fig. 4**), some of which include notorious pathogenicity-associated genera like *Clostridium*, *Bacillus*, *Enterococcus* or *Streptococcus* (**Table S2**). Functional investigations of the candidate RRNPP QSSs encoded by these genera could hence potentially unearth novel density-dependent mechanisms linked to pathogenicity. Moreover, this analysis reveals that some of the previously identified poly-encoding QSSs MGEs (**Table S3**) encode candidate RRNPP QSSs that belong to distinct groups/gene families (e.g. Rap and AimR in bacteriophages of *B. subtilis*; AloR13 and group5 in pBRLA3 plasmid of *B. laterosporus* (**Fig. 4**)) and could be thus considered as ‘bilingual’. From a host-MGE coevolution perspective, it is also interesting to note that besides the two lysis-lysogeny regulating AimR groups that should be 100% viral (Phaster failed to predict a prophage origin for 8 QSSs (unlike Prophage Hunter)), 6 groups of homologous QSSs were found to be shared between chromosomes and MGEs (**Fig. 3**). This suggests that some QSSs may be externalized between chromosomes, plasmids, and phages (45), with the possibility that such QSSs interfere with the regulation of genes in different genomes/ genetic elements. Typically, QSS families shared by MGEs and chromosomes can give rise to density-dependent manipulations of bacterial hosts by MGEs, as exemplified in our previous study on the host-derived Rap-Phr QSS of Bacillus phage phi3T (43) or in the study of Silpe and Bassler on the host-derived VqmA quorum sensing receptor of Vibrio phage VP882 (46).

**Figure 3:**
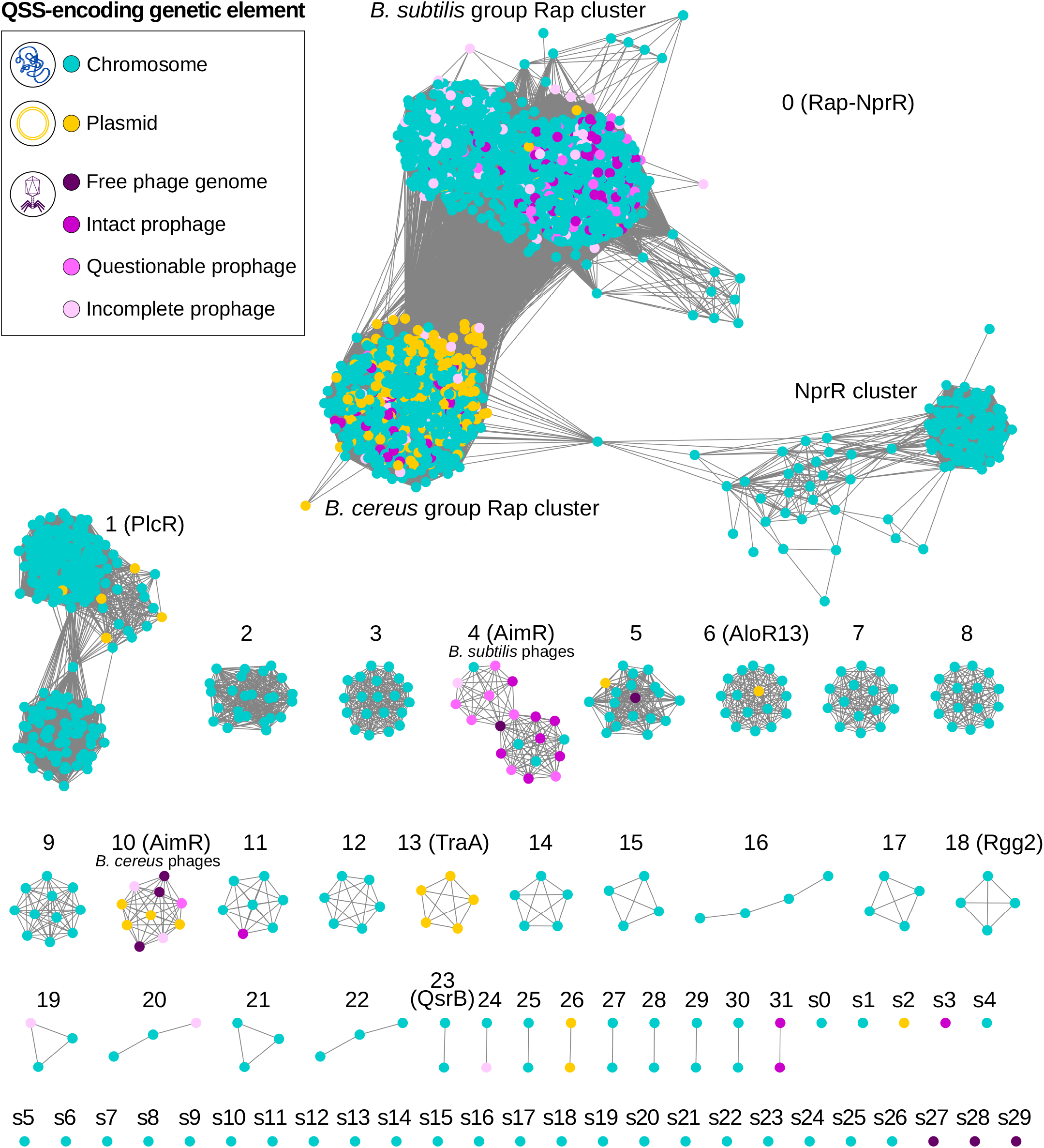
Sequence similarity network of the receptors forming a ‘conservative’ candidate RRNPP QSS with a cognate propeptide. Each node corresponds to a receptor sequence found adjacent to a candidate pro-peptide and is colored according to the type of genetic element encoding the QSS, as displayed in the legend. Each edge corresponds to a similarity link between two receptors defined according to the following thresholds: percentage identity >= 30%, alignment coverage >= 80% of the lengths of both receptors, E-value <= 1E-5. Each connected component of the graph thereby defines groups of homologous receptors, as in (44). The groups are ordered from the largest to the smallest. Labels that begin with an integer correspond to groups of homologous receptors (more than 1 sequence) whereas labels that begin with the ‘s’ character correspond to singletons (1 sequence only). A label followed by the name of a reference receptor in brackets means that at least one sequence in the group of homologous receptors is found to be homolog with a reference receptor according to the above thresholds.

**Figure 4:**
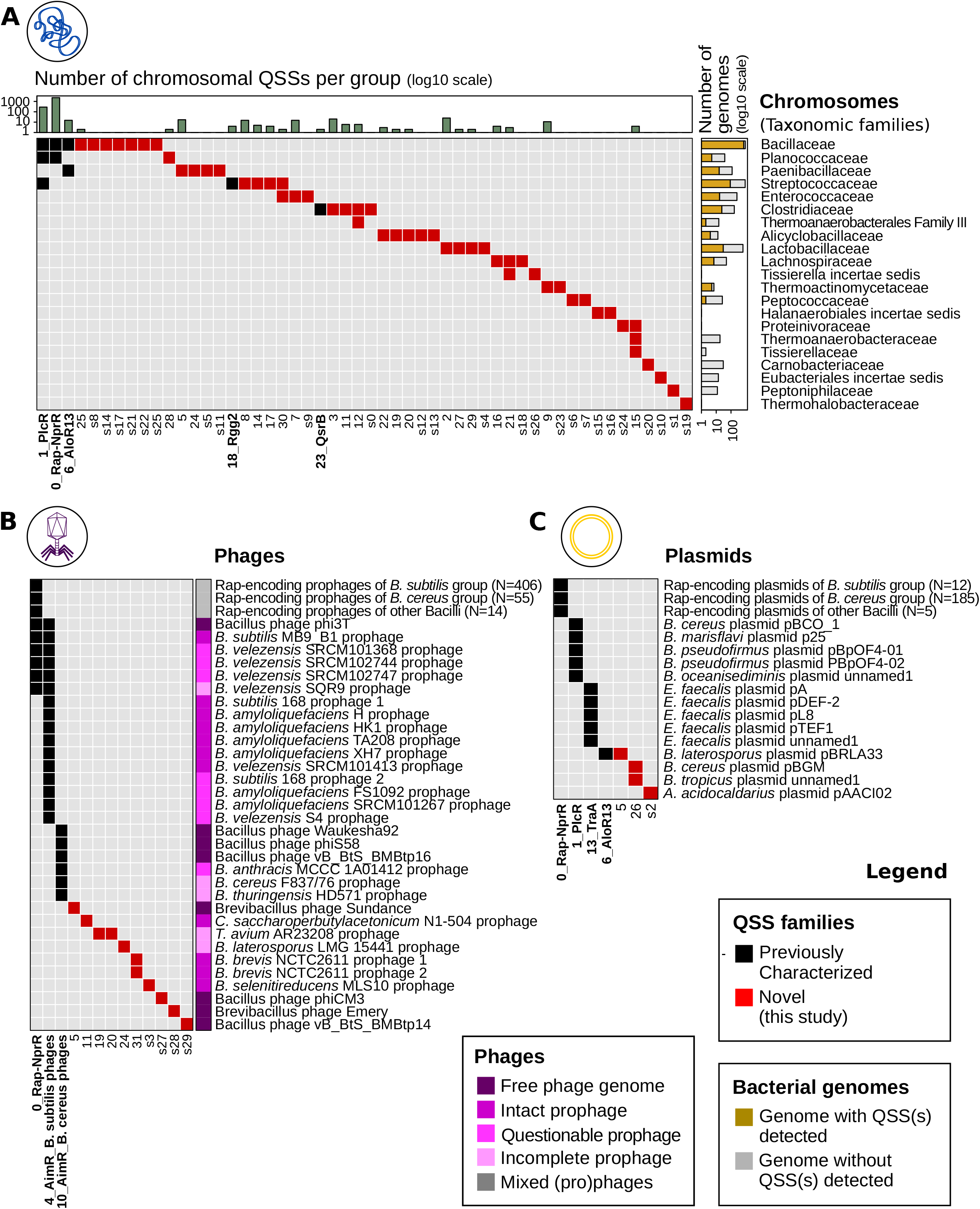
Distribution of groups of ‘conservative’ candidate RRNPP QSSs across taxonomic families (chromosomal QSSs) or MGEs (plasmidic and (pro)phage-encoded QSSs). **A.** Distribution of groups of RRNPP QSSs encoded by chromosomes (columns) across taxonomic families of the Firmicutes phylum (rows). Grey squares mean that no QSS of a given group has been detected in a given taxonomic family. Conversely, black and red squares mean that a given group has been detected in at least one chromosome of a given taxonomic family. Specifically, black squares highlight groups of RRNPP QSSs previously experimentally validated, whereas red squares represent candidate novel groups of RRNPP QSSs. The number of chromosomal representatives of each group of RRNPP QSSs is given in the histogram at the top of the heatmap (log10 scale). The number of chromosomes in each taxonomic family is given by the histogram at the right of the heatmap (log10scale): the gold area corresponds to the number of chromosomes in which at least one ‘conservative’ RRNPP QSS has been identified, whereas the grey area depicts the remaining chrosomomes without any RRNPP QSS detected. **B**. Distribution of groups of RRNPP QSSs across isolated genomes of temperate bacteriophages or prophages detected by Phaster within genomes of Firmicutes. Each row represents a single phage genome or a single prophage, to the exceptions of the first three rows that depict groups of prophages in which a single RRNPP QSS of the Rap family has been detected. The color of each (pro)phage indicates the quality of the prediction that the QSS-encoding entity is a phage: dark purple corresponds to sequenced phage genomes (100% sure), whereas lighter shades of purple correspond to different prophage qualities assessed by Phaster. **C**. Distribution of groups of RRNPP QSSs across plasmids sequenced in Firmicutes.

### Post-processing functional prediction of candidate RRNPP QSS reveals 50 putative QSS-regulated biosynthetic gene clusters

QSSs with a one component receptor module tend to be located nearby their target genes (47–49) and RRNPP QSSs are no exceptions to this trend, especially not MGE-encoded RRNPP QSSs (as shown in **Fig. 1**). The genomic neighborhood of a RRNPP QSS can thus provide insights on the potential function(s) regulated in a density-dependent by the QSS, as suggested in our previous study on the functional inference of viral RRNPP QSSs (https://www.biorxiv.org/content/10.1101/2021.07.15.452460v1). For instance, one function regulated by quorum sensing typically is the production of public good metabolites, *e.g*. antimicrobial compounds, because only a collective production may bring such molecules to the concentration levels required to exert a significant effect on the microbial community (50). Consistently, many natural product biosynthetic gene clusters (BGCs) have been demonstrated to be controlled by a QSS located in their genomic vicinity, be it a molecule-based QSS (48,49) or a peptide-based QSS (**Fig. 5A**) (24,25). To illustrate how the output of RRNPP_detector can be coupled with functional insights, we chose to focus on this practical example of putative QSS-regulated BGCs. Indeed, as a major challenge in the field of natural-product discovery is that many BGCs are not expressed under laboratory growth conditions (51), QSS-regulated BGCs are promising: they may represent a source of novel natural products for which the link to population density provides some understanding about how to elicit production. Hence, to identify candidate QSS-regulated BGCs, we first searched for BGCs with antiSMASH standalone version 6.0.0 (default parameters) (52) in the 474 genetic elements encoding at least one candidate RRNPP QSS with a receptor detected as a transcription factor (harboring a DNA binding domain). We then intersected the list of the 3153 BCGs detected by antiSMASH with our list of candidate RRNPP QSSs on the basis of the inclusion of the QSS region (from the start codon of the first gene to the stop codon of the second gene) within the region of a BGC defined by antiSMASH. This resulted in a subset of 46 candidate BGCs inferred to be regulated by RRNPP QSSs (**Table S4** and **Fig. 5**). Among these 46 putative QSS-regulated BGCs, 3 are plasmidic and are inferred by antiSMASH to produce antimicrobial peptides (**Fig. 5B**). As these plasmidic RRNPP QSSs likely enact the production of defense metabolites only when the quorum of plasmids is met, they might create a selective pressure for the acquisition of the plasmid by host cells at high plasmid densities, supporting complex scenarios of co-evolution. Interestingly, the 46 putative QSS-regulated BGCs produce major classes of natural products (**Table S4** and **Fig. 5C**), including ribosomally synthesized and posttranslationally-modified peptides (RiPPs), non-ribosomal peptides and polyketides. Worth to note, RiPPs are of most frequent occurrence, likely reflecting the important roles of RiPPs in bacterial physiology (53). Last but not least, the inference that these BGCs are expressed at high population densities hints at important ecological roles for their biosynthesized natural products, among which could be novel defense metabolites of medical relevance.

**Figure 5:**
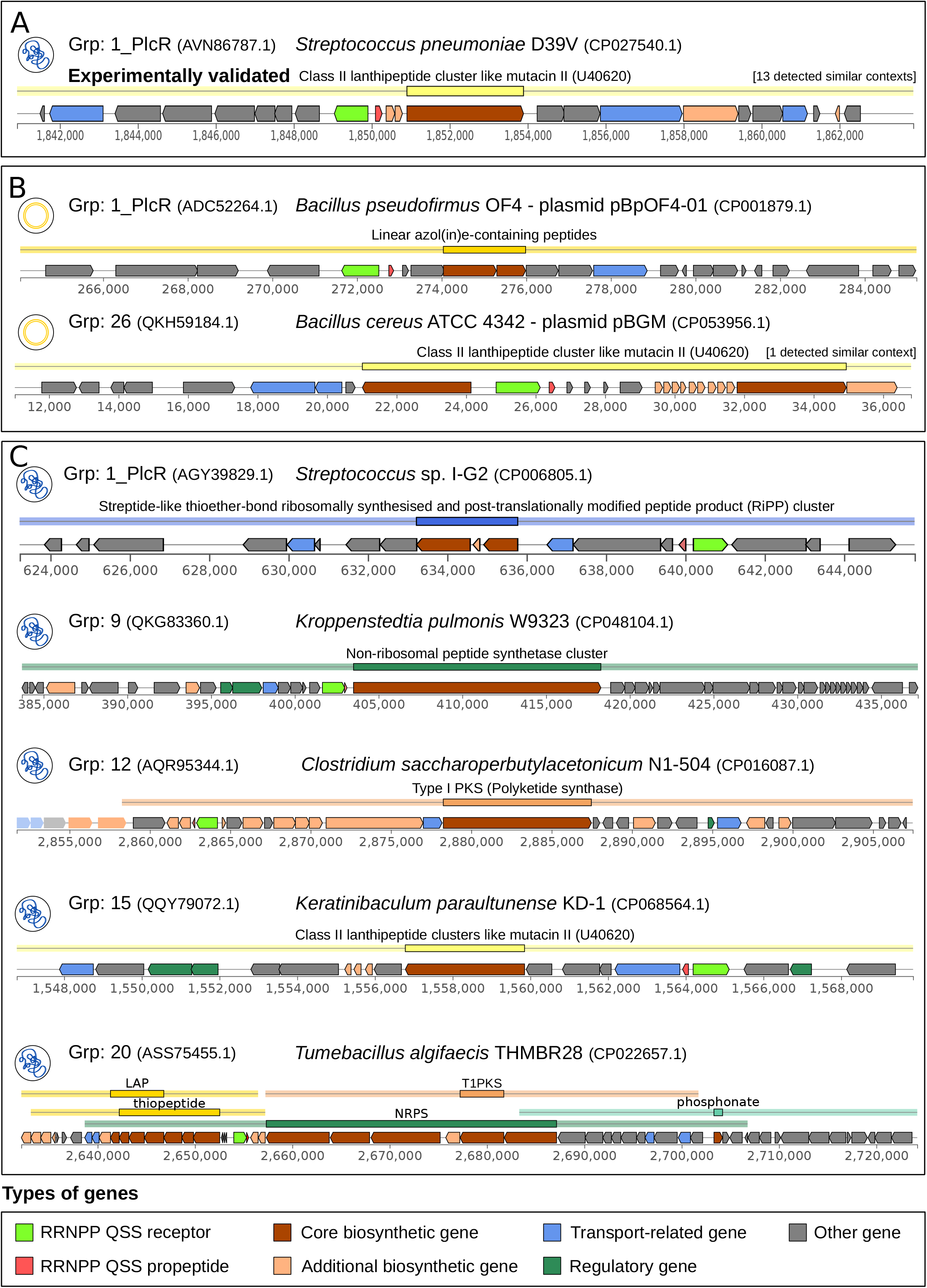
Selection of BGCs outputed by antiSMASH that are likely regulated by candidate RRNPP QSSs. Each genomic context corresponds to a BGC region outputed by antiSMASH within which a ‘conservative’ candidate RRNPP QSSs has been detected by RRNPP_detector. The genes are colored according to the legend displayed at the bottom of the figure. For each genomic context, a first line of information is composed of a thumbnail indicative of the genetic element encoding the BGC region (chromosome or plasmid). Then, a ‘receptor’ label indicates the group of homologous receptors to which the candidate RRNPP QSS receptor belongs and displays the NCBI ID of this receptor in brackets. Finally, the name of the genome along with its NCBI accession are given. The second line indicates the biosynthesis mode of the BGC as classified by antiSMASH, together with the number of similar genomic contexts identified in the taxon. The tickmarcks at the bottom of each BGC correspond to genomic coordinates, given in number of base pairs. **A.** Proof of concept: BGC demonstrated to be regulated by a RRNPP QSS in (24) and captured by our method. **B**. Plasmidic BGCs inferred to be regulated by a candidate RRNPP QSS. The context displayed for plasmid pBGM is conserved in plasmid unamed 1 of *B. tropicus*. **C**. Selection of chromosomal BGCs inferred to be regulated by candidate RRNPP QSSs to give an overview of putative QSS-regulated BGC diversity.

## CONCLUSION AND FUTURE PERSPECTIVES

RRNPP_detector is able to predict a wide range of candidate RRNPP QSSs in chromosomes or in mobile genetic elements (e.g. plasmids, phages) of gram-positive bacteria (e.g. Firmicutes). Because RRNPP_detector does not rely on sequence homology to known QSS components to detect candidate QSS gene families, it can detect novel families of RRNPP QSSs, as exemplified in this article. Moreover, as RRNPP QSSs tend to regulate adjacent genes in a density-dependent manner, post-processing genomic context analyses can couple RRNPP_detector output with functional insights, as shown in this article with the specific example of putative QSS-regulated BGCs.

RRNPP_detector presents a first step towards the large-scale identification of peptide-based QSSs. However, this software is dedicated to the sole detection RRNPP QSSs and we already anticipate to expand its usage to peptide-based QSSs with two-component systems. We believe that our work will unlock new biological knowledge regarding peptide-based biocommunication and will reveal many novel density-dependent decision-making processes in bacteria, plasmids and bacteriophages that could likely transform our views of microbial adaptation and bacteria-MGE co-evolution. Importantly, Firmicutes represent with Bacteroidetes the most prevalent phyla in human gut microbiomes (54), which makes the use of RRNPP_detector particularly relevant against human-associated metagenomics-assembled genomes or mobile genetic elements (e.g. from the human MGE database (55) or the Gut Phage Database (56)) to infer density-dependent behaviors that may take place within human intestinal microbiomes, plasmidomes and viromes.

## DATA AVAILABILITY

RRNPP_detector is freely available on github at **https://github.com/TeamAIRE/RRNPP_detector**

## SUPPLEMENTARY MATERIALS

- **Table S1:** 6467 ‘permissive’ candidate RRNPP QSSs
- **Table S2:** 3529 ‘conservative’ candidate RRNPP QSSs
- **Table S3:** 38 MGEs encoding multiple ‘conservative’ candidate RRNPP QSSs
- **Table S4:** 46 putative QSS-regulated BGCs

